# *Paenibacillus* sp. strain UY79, isolated from a root nodule of *Arachis villosa*, displays a broad spectrum of antifungal activity

**DOI:** 10.1101/2021.03.26.437297

**Authors:** Andrés Costa, Belén Corallo, Vanesa Amarelle, Silvina Stewart, Dinorah Pan, Susana Tiscornia, Elena Fabiano

## Abstract

A nodule-inhabiting *Paenibacillus* sp. strain (UY79) isolated from wild peanut (*Arachis villosa*) was screened for its antagonistic activity against diverse fungi and oomycetes (*Botrytis cinerea, Fusarium verticillioides, Fusarium oxysporum, Fusarium graminearum, Fusarium semitectum, Macrophomina phaseolina, Phomopsis longicolla, Pythium ultimum, Phytophthora sojae, Rhizoctonia solani, Sclerotium rolfsii* and *Trichoderma atroviride*). Results obtained show that *Paenibacillus* sp. UY79 was able to antagonize these fungi/oomycetes and that agar-diffusible metabolites and enzymes, as well as volatile compounds (different from HCN), participate in the antagonism exerted. We found that *Paenibacillus* sp. strain UY79 did not affect symbiotic association or growth promotion of alfalfa plants when co-inoculated with rhizobia. By whole genome sequence analysis, we determined that strain UY79 is a new species of *Paenibacillus* within the *Paenibacillus polymyxa* complex. Diverse genes putatively involved in biocontrol activity were identified in the UY79 genome. Moreover, according to genome mining and antibiosis assays, strain UY79 would have the capability to modulate the growth of bacteria commonly found in soil/plant communities.

**IMPORTANCE:** Phytopathogenic fungi and oomycetes are responsible for causing devastating losses in agricultural crops. Therefore, there is an enormous interest in the development of effective and complementary strategies that allow the control of the phytopathogens, reducing the input of agrochemicals in croplands. Discovery of new strains with expanded antifungal activities and with a broad spectrum of action is challenging and of great future impact. Diverse strains belonging to the *P. polymyxa* complex have been reported to be effective biocontrol agents. Results presented here show that the novel discovered strain of *Paenibacillus* sp. presents diverse traits involved in antagonistic activity against a broad spectrum of pathogens and would be a potential and valuable strain to be further assessed for the development of biofungicides.

## INTRODUCTION

Plant health depends on the existence of a proper balance between beneficial and pathogenic microorganisms with which plants coexist in natural environments. In unbalanced environments, pathogenic fungi and oomycetes may produce severe plant diseases. It has been estimated that crop losses due to fungal diseases are around 30 % of world agricultural production (1). In healthy environments, various microorganisms able to suppress plant diseases, either by boosting plant immune system or by direct inhibition of the pathogen, could be found (2). Microorganisms with the ability to control the development of phytopathogens belong to the group of Biological Control Agents (BCAs), defined as “a natural enemy, antagonist, or other organism, used for pest control” (ISPM 05, International Standards for Phytosanitary Measures) (3).

Several mechanisms involved in the biocontrol of phytopahogens have been found in BCAs, ranging from hyperparasitism and predation to the production of lytic enzymes such as chitinases, cellulases, glucanases, proteases and lipases, which can disrupt the cell walls of many phytopathogenic fungi/oomycetes (4). Secretion of antibiotics or the generation of organic as well as inorganic volatile compounds, are also widespread biocontrol mechanisms reported (5-7). Successful BCAs generally express multiple biological traits that act additively and synergistically to efficiently suppress the pathogen (6). BCAs are ubiquitous constituents of soils and also of plant microbiota. They can be found as endophytes of roots, steams, leaves and flowers of diverse plant genera, and also in legume nodules (8-10). A remarkable feature of legumes is their ability to establish symbiotic associations with a group of bacteria known as rhizobia, belonging to the orders Rhizobiales and Burkholderiales. As part of this symbiosis, distinctive structures, called nodules, are elicited in the root (or occasionally in the stems) of the legume where the biological nitrogen fixation process takes place (11). Legume nodules were considered for many years to be exclusively occupied by rhizobia, however strong evidences demonstrated that nodules also harbor a diverse community of microorganisms (10, 12-14). Major bacterial phyla that have been consistently found as nodules inhabitants include *Actinobacteria, Firmicutes*, and *Proteobacteria*. The presence of *Agrobacterium, Arthrobacter, Acinetobacter, Bacillus, Bosea, Enterobacter, Micromonospora, Mycobacterium, Paenibacillus, Pseudomonas*, and *Stenotrophomonas* genera were reported, being strains of *Bacillus* spp. and *Paenibacillus* spp. frequently isolated from this niche (10, 12, 13). The role of nodule endophytes is unknown, but it has been found that some strains possess biocontrol activity against phytopathogens (14). This fact makes nodules an interesting and still poorly explored environment for the identification of strains with biocontrol activity against phytopathogenic fungi.

The goal of this work was to identify and characterize a nodule-inhabiting bacterium obtained from *Arachis villosa* (wild peanut) collected in a Uruguayan National Park. According to the genome analysis, strain UY79 is a new species of *Paenibacillus* that belongs to the *Paenibacillus polymyxa* group. Results obtained *in vitro* and *in silico* indicate that strain UY79 possesses a broad-spectrum of antimicrobial activity, as well as a plethora of mechanisms putatively involved in antagonisms.

## MATERIALS AND METHODS

### Microbes, media and growth conditions used in this work

Microorganisms used in this work are listed in Table S1. The bacterial strain UY79 was isolated from a root nodule of *A. villosa*, a native Uruguayan legume, collected in Nuevo Berlin, Río Negro, Uruguay (S32°59’04.00” W58°03’48.20”). The nodule was surface sterilized for 2 min with a solution of 10 mM HgCl_2_ in 0.1N HCl, followed by seven washes with sterile distilled water. The surface sterilized nodule was crushed with a sterile glass rod and streaked on Yeast-Mannitol Agar (15) supplemented with 1 g/l glutamate (YMAG). Plates were incubated at 30 °C and checked every day for bacterial growth. A colony phenotypically different from rhizobia was selected, cultured in YMG broth and stored at - 80 °C with 25 % (v/v) glycerol.

Strain UY79 was then routinely grown in Tryptic Soy Broth/Agar (TSB/TSA, BD™) or Potato Dextose Broth/Agar (PDB/PDA, Oxoid Ltd.) at either 30 °C or 25 °C, as indicated throughout the manuscript. Fungi and oomycetes were routinely grown on PDA at 25 °C, except for *Phytophthora sojae* Ps25 that was grown on V8 agar medium (16). Strains of fungi and oomycetes used in this work were obtained from two culture collections: Laboratorio de Micología, Facultad de Ciencias and Instituto Nacional de Investigación Agropecuaria, La Estanzuela.

### Phylogenetic affiliation of strain UY79 using *16S rRNA*

Genomic DNA was purified using the commercial kit Zymo Quick-DNA™ Fungal/Bacterial Miniprep Kit as described by the manufacturer.

An almost complete sequence (c.a. 1,400 bp) of the *16S rRNA* gene was obtained by PCR amplification using the universal primers 27F (5′-AGAGTTTGATCMTGGCTCAG-3′) and 1492R (5′-TACGGYTACCTTGTTACGACTT-3′) (17) as previously described (18). Amplicons were sequenced at Macrogen Inc. (Seoul, Korea). Forward and reverse sequences were assembled and curated using the DNA Baser V3 Sequence Assembler. Sequence obtained was deposited in the NCBI GenBank database with the Accession Number MT973969. Identification of bacterial genus was accomplished using the “Identify” tool at the EzBioCloud server (19) (https://www.ezbiocloud.net/identify).

In order to evaluate if strain UY79 was phylogenetically related to other *Paenibacillus* strains isolated from similar environments, a phylogenetic tree was constructed including *16S rRNA* gene sequences from strain UY79, thirty-three *Paenibacillus* spp. type strains isolated from soil, compost, rhizosphere, or different plant compartments and tissues (root nodule, root, leaf, seed), as well as two pathogenic strains (*Paenibacillus larvae* ATCC 9545 and *Paenibacillus lautus* NBRC 15380). Sequences were retrieved from EzBioCloud. Sequences were aligned with MAFFT v7.453, and Gblock v0.91b was used to remove poorly aligned positions. A Maximum Likelihood tree was assembled in MEGA X based on the Kimura 2-parameter model (+G+I) (20, 21). Robustness of the tree branches was estimated with 10,000 bootstrap pseudoreplicates.

### Sequencing, assembly, annotation and mining of *Paenibacillus* sp. UY79 genome

The genome of strain UY79 was sequenced by paired-end-sequencing method using Illumina TrueSeq platform (Macrogen, Seoul, Korea). Low quality sequences were removed using Trim Galore 0.4.4 with the following line command: trim_galore --paired --three_prime_clip_R1 10 --three_prime_clip_R2 10 --length 50 UY79_1.fastq UY79_2.fastq. Filtered sequences were used for de novo assembly using Velvet Assembler 2.2.5 with the following line command: VelvetOptimiser.pl -s 19 -e 81 -d AssemVelvetOptimizer -f ‘-shortPaired -fastq -separate UY79_1_val_1.fq UY79_2_val_2.fq’ -t 8 --. The genome was annotated by using the NBCI Prokaryotic Genome Annotation Pipeline (22) and the Rapid Annotation using Subsystem Technology (RAST 2.0) (23). Strain UY79 genome sequence obtained was deposited in the NCBI GenBank database with the Accession Number SUB8965727. In order to identify gene clusters putatively encoding for secondary metabolites with antimicrobial activity we used the web server ANTISMASH 5.2.0 based on profile hidden Markov models of specific genes (24). The PROPHAGE HUNTER and PHASTER web servers were used to predict and annotate bacteriophage genes in the bacterial genome (25, 26). BLAST tool (27) on NCBI web server was used for manual annotation of some genes.

### Multilocus sequence analysis and average nucleotide identity test

A phylogenetic analysis based on multilocus sequences (MLSA) was performed using *16S rRNA, gyrB, rpoB, recA* and *recN* concatenated gene sequences (8,535 positions) retrieved from the sequenced *Paenibacillus* spp. genomes publicly available at EzBioCloud database http://www.ezbiocloud.net/eztaxon (19). Sequences were aligned with MAFFT v7.453, and Gblock v0.91b was used to remove poorly aligned positions. A Maximum Likelihood tree was assembled in MEGA X, based on the Maximum likelihood method and General Time-Reversible model (+G+I)(28). Robustness of the tree branches was estimated with 1,000 bootstrap pseudoreplicates.

Average Nucleotide Identity (ANI) score between the species included in the MLSA was calculated using the ANI calculator tool from EzBioCloud (19) (https://www.ezbiocloud.net/tools/ani).

### *In vitro* antagonistic activity of strain UY79 against fungi and oomycetes

For antagonism assays, the strain UY79 was grown in 5 ml of TSB for 16 h at 30 °C and 200 rpm. *Macrophomina phaseolina* J431, *Rhizoctonia solani* Rz01, *Pythium ultimum* Py03, *Fusarium graminearum* S127, *Fusarium oxysporum* J38, *Phomopsis longicolla* J429, *Sclerotium rolfsii* 1948, *Fusarium verticillioides* A71, *Fusarium semitectum* J41, *Botrytis cinerea* A1 and *Trichoderma atroviride* 1607 were cultured on PDA and *Phytophthora sojae* Ps25 was cultured on V8 agar medium, at 25 °C for 5 days.

To evaluate the antifungal activity of agar-diffusible compounds produced by strain UY79, a dual plate assay was performed as described by Geels and Schippers (29). Briefly, a 0.9 cm mycelial agar-plug from the leading edge of the fungus/oomycetes culture, previously grown for 5 days at 25 °C, was placed on a fresh PDA medium. Strain UY79 was streaked as a small line approximately 3 cm away from the mycelial plug. Plates were incubated at 25 °C for 2-8 days, until radial growth of the mycelia reached the edge of the plate opposite to the side with bacteria inoculum. Mycelial growth diameter and formation of inhibition zones around bacterial growth was recorded. Three independent assays were performed.

In order to detect the presence of antifungal compounds in bacterial cultures grown in liquid medium, the strain UY79 was grown in 100 ml of PDB medium at 25 °C and 200 rpm. After 30-, 44-, 68-, 88- and 96 hours, 10 ml were collected, centrifuged at 10,000xg for 10 min, and supernatants were filtered through a 0.45 µm filter. Cell-free supernatants were mixed with melted (45 °C) PDA (40 g/l agar) medium (1:1) and 2 ml of the mixture was poured into a 6-well plate. A 0.5 cm mycelial plug from a *F. verticillioides* fresh culture was placed in the center of the well, incubated 3 days at 25 °C and mycelial growth was measured. Fungal growth on PDA medium without the addition of supernatant was used as a control. Three independent assays were performed.

To evaluate the production of volatile compounds (VCs) with antifungal activity, a dual plate assay (30) was conducted in PDA or V8 agar as indicated. Briefly, strain UY79 was grown in 5 ml of TSB for 16h at 30°C and 200 rpm, and 100 µl of the culture was spread as a lawn in a Petri dish. On a different Petri dish, a 0.9 cm mycelial plug of the fungi/oomycetes was placed in the center. Both uncovered Petri dishes were faced to each other, and sealed with parafilm to prevent VCs leakage. As controls, Petri dishes containing the fungi/oomycetes were faced to plates with PDA or V8 agar medium as indicated. Plates were incubated at 25 °C until the mycelia of the control plates reached the edge of the Petri dish. Antagonism was determined by measuring the percentage of growth inhibition (GI %) as follows: 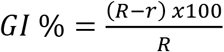 where R is the radio of the mycelia of the control fungus/oomycete not faced to the bacterium, and r is the radio of the fungus/oomycetes faced to the bacterium. Three independent biological replicates were performed.

Production of the volatile compound hydrogen cyanide was assessed qualitatively by the picrate-filter paper method (31). A dual plate assay was performed, where picrate embedded filter paper was placed in the cover of a Petri dish, face to face with a lawn of bacteria. The plate was sealed with parafilm to prevent VCs leakage. Both PDA and TSA medium, either with the addition of 4.4 g/l of glycine or in its absence were assessed. Cyanogenic activity was visualized as a change in color of the filter paper from yellow to orange. *Pseudomonas fluorescens* UP61 was used as a positive control and medium without bacteria as a negative control.

### Determination of siderophore production, and xylanase, β-glucosidase, cellulase and protease activities

Strain UY79 was grown in TSB for 48 h at 30 °C and 200 rpm, and 10 µl drops were spotted in each of the media to be assayed. Three independent assays were performed in all the experiments.

Siderophore production was assessed as the formation of an orange halo around the colony, by using the chromo azurol sulfonate (CAS) agar plate method (32). To evaluate xylanase activity, the bacterium was grown on TSA containing 0.5 % (w/v) xylan beechwood as a substrate. Xylanase activity was visualized as a clear halo around the colony (33).

β-glucosidase activity was assessed on TSA containing 0.2 % (w/v) esculin and 0.03 % (w/v) FeCl_3_. The esculetin released from esculin by β-glucosidase action was detected as a dark halo around the colonies (34). *Escherichia coli* DH5α β glu and *Pseudomonas putida* KT2440 β glu were used as positive controls while *E. coli* DH5α and *P. putida* KT2440 as negative controls.

Cellulolytic activity was assayed on bacterial cultures grown on TSA containing 0.5 % (w/v) of carboxymethyl cellulose. Cellulase activity was detected using congo red staining (35). Briefly, colonies were overlaid with 0.05 % (w/v) congo red, incubated for 10 min at room temperature, washed with distilled water, and incubated for 10 min with 1 M NaCl. An orange halo around colonies in the red background indicates cellulose-hydrolyzing activity. *E. coli* DH5α cel5 and *P. putida* KT2440 cel5 were used as positive controls while *E. coli* DH5α and *P. putida* KT2440 as negative controls.

Proteolytic activity was assessed on bacterial cultures grown on skim milk agar. A solution of autoclaved skim milk 10 % (w/v) was added in 1:1 (v/v) ratio to TSA diluted medium 1: 100 to achieve a final concentration of 5 % of skim milk. Protease activity was detected as a clear halo around the colony (36).

### *In vitro* evaluation of direct plant growth promoting activities

In order to determine if strain UY79 has the potential to fix nitrogen, the presence of *nifH* gene (a gene that encodes the nitrogenase iron protein) was assessed by PCR using the PolF (5’-TGCGAYCCSAARGCBGACTC-3’) and PolR (5’-ATSGCCATCATYTCRCCGGA-3’) primers, which amplify a 350 bp intragenic region (37). *Ensifer meliloti* 1021 was used as a positive control.

Indole-3-acetic acid (IAA) production by strain UY79 was determined by a colorimetric assay using the Salkowski reagent according to the method described by Gordon and Weber (38). Briefly, an IAA-producing strain (*Microbacterium sp*. strain UYFA 68) and the strain UY79 were grown in 6 ml of TSB with or without 100 µg/ml tryptophan. The IAA concentration in cultures was determined from a calibration curve performed with pure IAA. Each treatment was performed thrice.

To evaluate the ability to solubilize phosphate, bacteria were grown in NBRIP (39) or PVK (40) media, containing rock phosphate or (PO_4_)_2_Ca_3_, as the sole phosphate source. Solubilization of phosphate was visualized as a clear halo around the colony. *Pantoea* sp. UYSB45 was used as a positive control. Three independent assays were performed.

### Effect of strain UY79 in alfalfa plant growth promotion exerted by two rhizobia strains

Seeds of *Medicago sativa* var. crioula were surface sterilized with 10 mM HgCl_2_ in 0.1N HCl as previously described (41). Surface-sterilized seeds were germinated at 30 °C on Petri dishes containing water with 0.8 % (w/v) agar. After germination, seedlings were transferred into glass plant tubes containing 15 ml of Jensen’s N-free medium (15) with 0.8 % (w/v) agar. Seedlings were inoculated simultaneously with *Paenibacillus* sp. strain UY79 and either *E. meliloti* 242 or *E. meliloti* 1021. Inoculation was done using c.a. 1×10^7^ CFU of each strain. The experiment included the following controls: plants solely inoculated with either *E. meliloti* 242, *E. meliloti* 1021, or with strain UY79. A control without bacteria was included. Plants were grown at 24 °C with a photoperiod of 16 h light and 8 h darkness. Two independent assays were performed and plants were harvested after 6- or 10-weeks post-inoculation (assays I and II, respectively). Dry weight of the aerial portion and nodule number per plant were recorded. Eight plants (one plant per tube) were used per conditions. Experiments were independently analyzed using the nonparametric Kruskal-Wallis test and pairwise comparisons using Wilcoxon test for dry weight and an ANOVA with post-hoc Tukey HSD test for nodule count. An alpha of 0.05 was used as the significance cut-off value for all statistical analyze.

### Assessment of antibiosis activity against different soil and plant associated bacteria

To investigate if strain UY79 displayed antibacterial activity, eleven gram-negative strains (belonging to the *Bradyrhizobium, Ensifer, Rhizobium, Cupriavidus, Paraburkholderia, Azospirillum* and *Erwinia* genera) and two gram-positive strains (*B. subtillis* ATCC 6633 and *Streptomyces* sp. UYFA156) were used as target strains (Table S1). The soft-agar overlay assay was carried out according to James *et al*. (42) and Rao *et al*. (43) with slight modifications. Briefly, target strains were grown until late exponential phase in either TSB or TY media and 50 µl of the bacterial culture were used to inoculate 25 ml of soft agar (45 °C), and poured into either TSA or TY plates, respectively. A 10 µl drop of a UY79 culture grown in TSB for 24 h at 30 °C was spotted onto the inoculated solidified soft agar, and plates were incubated at 30 °C. Antibiosis was considered positive when a zone of inhibition, around or into the spots containing the UY79 inoculum, was observed. The assay was performed twice.

## RESULTS

### Identification of strain UY79, genome assembly and annotation

In this work we isolated a colony from a surface sterilized nodule of *A. villosa* with phenotypic traits different from that expected for rhizobia and characteristic of *Paenibacillus*/*Bacillus*. The isolate, named as UY79, was preliminary identified according to 16S *rRNA* sequence analysis using the EzBioCloud server. The three most similar sequences corresponded to *Paenibacillus ottowii* MS2379(T), *Paenibacillus peoriae* KCTC 3763(T) and *Paenibacillus polymyxa* NCTC 10343(T), with sequence identity of 99.41 %, 99.04 % and 98.96 %, respectively. Considering the fact that many *Paenibacilllus* strains have been isolated from plants or from plant-associated environments, we assessed if there was a phylogenetic association between phylogenetic position and the environmental origin of the strains. As shown in Fig. S1, no such correlation was observed. Moreover, *Paenibacillus kribbensis, P. peoriae, Paenibacillus jamilae* (recently reclassified as *P. polymyxa*)(44), *P. ottowii, Paenibacillus terrae, Paenibacillus brasilensis* and *P. polymyxa* (referred to as the *Paenibacillus polymyxa* complex) formed a monophyletic group, which is in agreement with previous reports (45), and strain UY79 clearly grouped within the *P. polymyxa* complex (Fig. S1).

As the resolution of 16S *rRNA* phylogeny is not enough to discriminate between *Paenibacillus* species, we sequenced the UY79 genome and performed a MLSA analysis based on 16S *rRNA, gyrB, rpoB, recA* and *recN* genes concatenated sequences retrieved from the genomes. As shown in Fig. 1, strain UY79 grouped close to *P. kribbensis, P. brasilensis* and *P. peoriae* type strains but in a different branch with a 100 % support based on 10,000 bootstrap pseudoreplicates. It is also interesting to notice that *P. polymyxa* strains grouped in three well supported clusters, as has been previously reported by Jeon *et al*. (46). Comparing the strain UY79 genome to the twenty-two sequenced genomes of the *Paenibacillus* species publicly available in GenBank, we found that values of average nucleotide identity (ANI) were below 88.9 % in all cases, indicating that the strain UY79 is a new *Paenibacillus* species.

**Fig 1.**
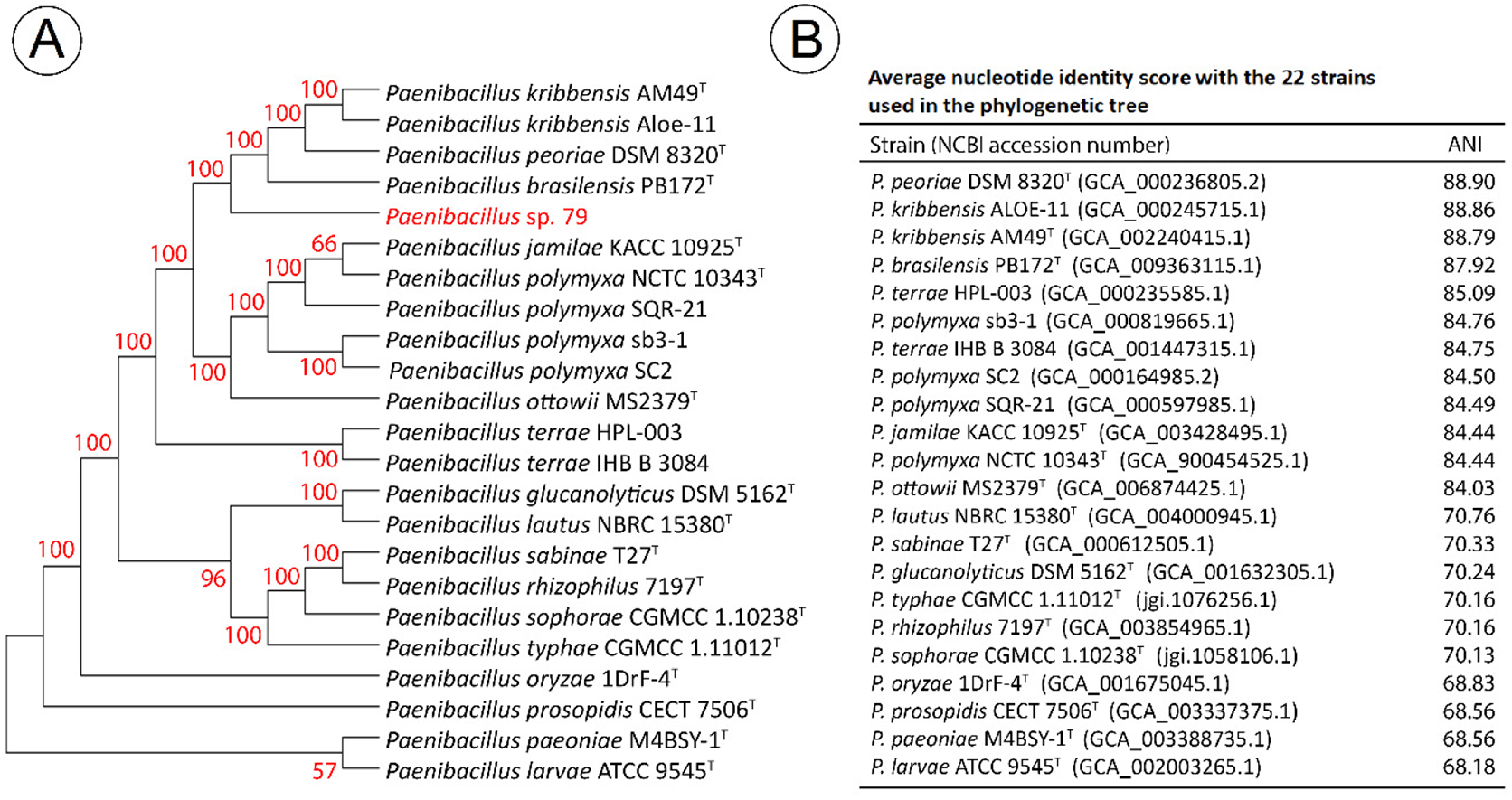
Taxonomic affiliation of *Paenibacillus* sp. strain UY79. **A**) Multilocus phylogenetic analysis based on 16S *rRNA, gyrB, rpoB, recA* and *recN* genes concatenated sequences (8535 positions) obtained from sequenced genomes. The tree was constructed using MEGAX program, Maximum likelihood method and General Time-Reversible model. Numbers at each node represent percentage of bootstrap replications calculated from 1,000 replicate trees. Strain UY79 is shown in red print, type strains are indicated with a superscript T. **B)** Scores of average nucleotide identity (ANI). NCBI accession numbers of genomes are shown in parentheses.

UY79 genome was assembled in 167 contigs with 235.7X average genome coverage. Genome size was calculated to be 4.9 Mb with 5353 predicted coding sequences according to RAST 2.0, and GC content was estimated to be 46.4 % (Table S2). A great variation has been found in genome sizes and GC content of *Paenibacillus* strains, with values ranging from 4 to 8.8 Mb for genome sizes and from 41 to 63 % in GC content (47, 48). Therefore, both genome size and CG content of UY79 genome are within the lowest ranges found.

### Strain UY79 exhibited a broad spectrum of antagonism against fungi and oomycetes

The *Paenibacillus* genus is well reported as antagonist of phytopathogenic fungi, being this effect mediated by different mechanisms which might involve the production of volatile or diffusible compounds with antimicrobial activity, the synthesis of cell wall degrading enzymes, and production of iron scavengers, among others (45, 46). First, we investigated if strain UY79 produced diffusible compounds with antagonistic activity and found that it was able to inhibit all ten fungi and the two oomycetes tested (Fig. 2). These results indicate that strain UY79 produced diffusible compounds in PDA (or V8 agar in the case of *P. sojae*) medium with a wide spectrum of antifungal activity. Although no quantitative evaluation was performed, qualitatively we can infer that strain UY79 produce an important mycelial growth inhibition of *B. cinerea* A1 (Fig. 2a), *F. verticillioides* A71 (Fig. 2e), *M. phaseolina* J431 (Fig. 2f), *P. longicolla* J429 (Fig. 2g), *P. sojae* Ps25 (Fig. 2h) and *R. solani* Rz01 (Fig. 2j); a moderate growth inhibition of *F. graminearum* (Fig. 2b), *P. ultimum* Py03 (Fig. 2i) and *T. atroviride* 1607 (Fig. 2l); and a slight growth inhibition of *F. oxysporum* J38 (Fig. 2c), *F. semitectum* J141 (Fig. 2d) and *S. rolfsii* 1948 (Fig. 2k). In addition, for *F. verticillioides* A71 faced to UY79, a zone of mycelial lysis at the edge of the mycelial growth facing the bacterium was observed (Fig. 2e). In the case of *R. solani* Rz01 (Fig. 2j) and *S. rolfsii* 1948 (Fig. 2k) a lower density of resistance structures was detected in the proximity of the confrontation zone. Moreover, it is interesting to notice the dark brown pigmentation of strain UY79, when faced to *R. solani* Rz01 (Fig. 2j).

**Fig 2.**
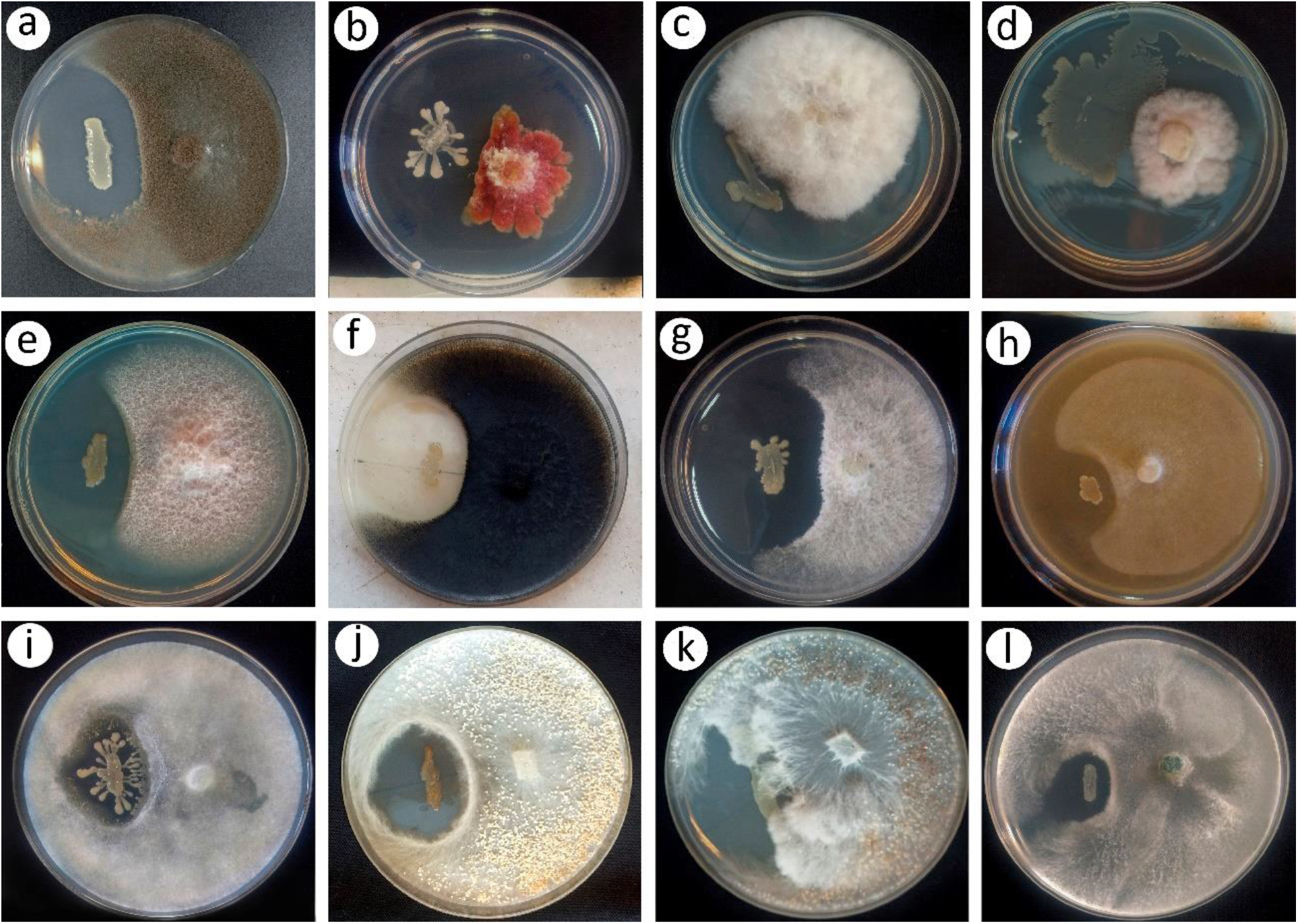
*In vitro* antagonistic activity of *Paenibacillus* sp. strain UY79 against fungi and oomycetes analyzed by the dual plate assay. Antagonistic activity of strain UY79 (left strike) was visualized as a growth inhibition zone of the fungi/oomycetes. *Botrytis cinerea* A1 (**a**), *Fusarium graminearum* S127 (**b**), *Fusarium oxysporum* J38 (**c**), *Fusarium semitectum* J41 (**d**), *Fusarium verticillioides* A71 (**e**), *Macrophomina phaseolina* J431 (**f**), *Phomopsis longicolla* J429 (**g**), *Phytophthora sojae* Ps25 (**h**), *Pythium ultimum* Py03 (**i**), *Rhizoctonia solani* Rz01 (**j**), *Sclerotium rolfsii* 1948 (**k**) and *Trichoderma atroviride* 1607 (**l**). Dual plate assays were performed on PDA medium, with the exception of *P. sojae* Ps25 that was grown on V8 agar medium. All assays were performed in triplicate and representative plates are shown.

As strain UY79 proved to be capable of producing diffusible compounds with antimicrobial activity in the presence of the fungi/oomycetes, we wondered if it was also able to produce them when grown alone in liquid medium. *F. verticilloides* A71 was randomly selected as target for this assay. As shown in Fig. 3, compounds that inhibited *F. verticillioides* A71 growth were present in cell-free supernatant of UY79 cultures after 30 h of incubation. The inhibition phenotype was maintained in 44-, 68-, 88- and 96 h cultures, suggesting that either antifungal compound/s production was maintained for up to 96 h, or that the produced compounds were active for such a period.

**Fig 3.**
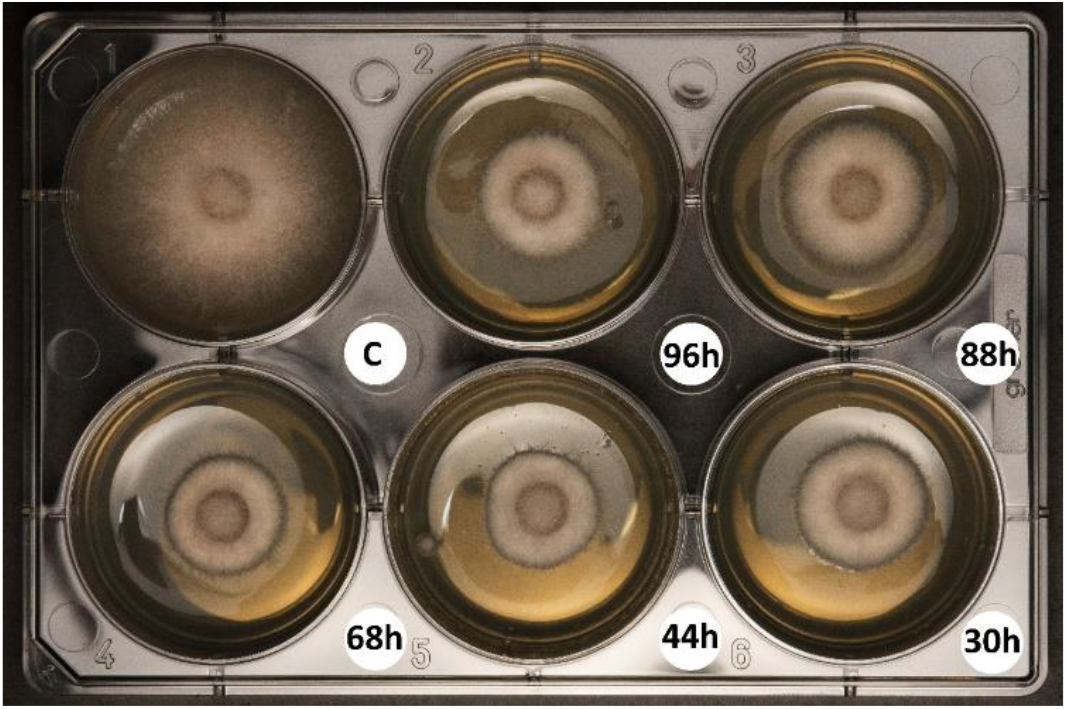
Cell-free supernatants of *Paenibacillus* sp. strain UY79 exhibit antifungal activity against *Fusarium verticillioides* A71. Cell-free supernatants of cultures grown for 30-, 44-, 68-, 88- and 96 h were included in PDA medium (1:1). PDB medium was used as control (C). Experiment was performed in triplicate and a representative plate is shown.

Next, we evaluated the capability of *Paenibacillus* sp. strain UY79 to produce volatile compounds (VCs) with antagonistic activity. As shown in Fig. 4A, VCs produced by strain UY79 were capable of inhibiting nine out of ten fungi, and the two oomycetes analyzed. Growth of seven phytopathogens, out of the eleven assessed, was inhibited more than 50 % (Fig. 4B). This was the case of *B. cinerea* A1 (a), *F. graminearum* S127 (b), *F. semitectum* J41 (d), *P. longicolla* J429 (g), *P. sojae* Ps25 (h), *R. solani* Rz01 (j) and *S. rolfsii* 1948 (k) (Fig. 4A and B). Remarkably, growth of *R. solani* Rz01 (j) and *B. cinerea* A1(a) was severely compromised (93±6 % and 72±11 %, respectively) when faced to strain UY79. *M. phaseolina* J431 (f) and *F. verticilliodes* A71 (e) were the least affected, resulting in an approximately 30 % of mycelial inhibition by UY79 VCs. As shown in Fig. 4A, an altered pigmentation was observed in *F. graminearum* S127 (b) and *M. phaseolina* J431 (f), and to a lesser extent in *F. semitectum* J41 (d) and *F. verticillioides* A71 (e), as a response to VCs produced by strain UY79. Moreover, lower density of hyphae was observed in *R. solani* Rz01 (j) while, on the contrary, a higher density of hyphae was produced by *P. sojae* Ps25 (h). Although *T. atroviride* 1607 (l) did not show a radial growth inhibition, an alteration in the colony pattern was observed. Altogether, these results indicate that strain UY79 is able to produce VCs with a broad spectrum of antimicrobial activity and that fungi/oomycetes responses depend on each particular microorganism.

**Fig 4.**
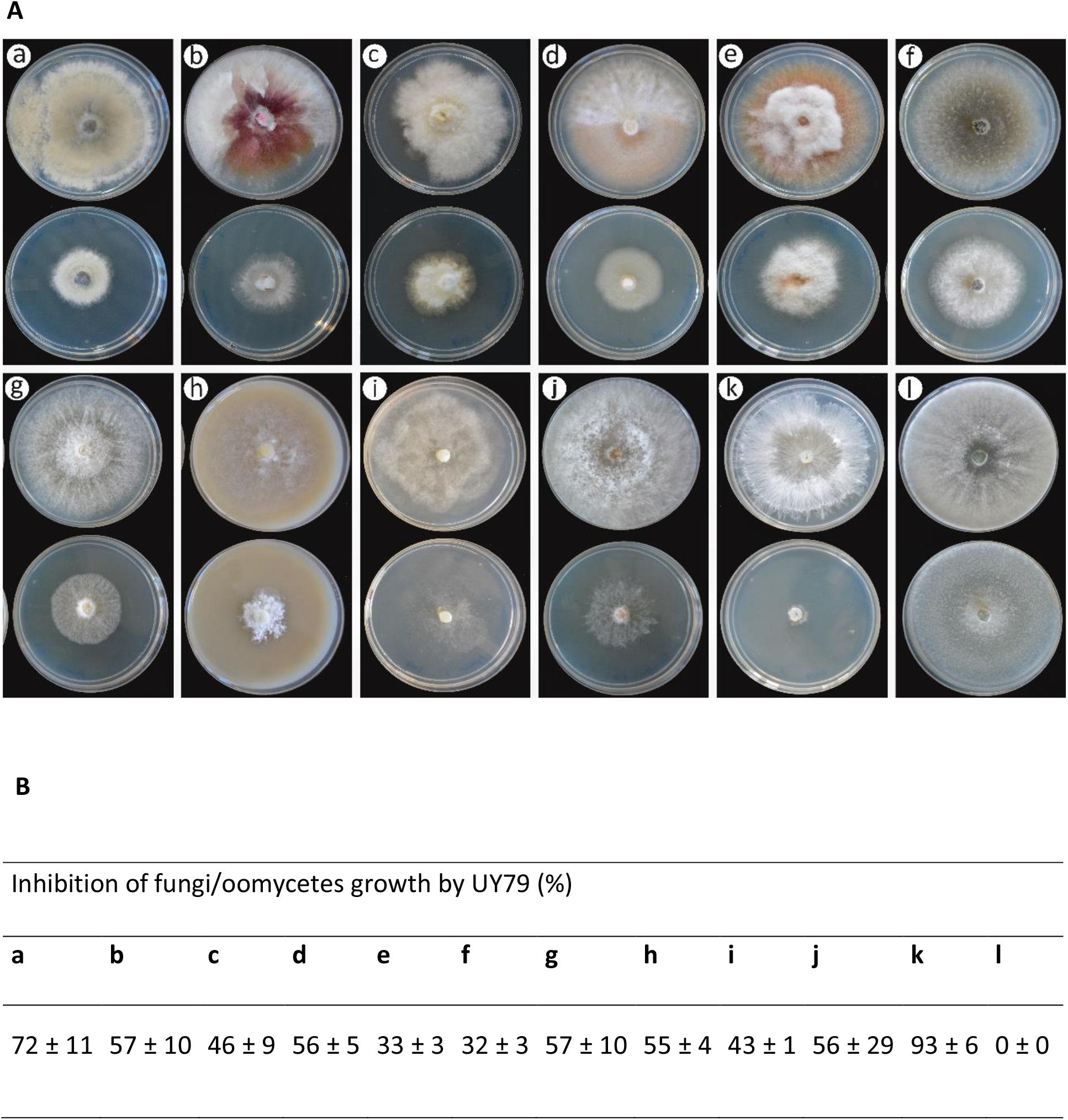
Volatile compounds produced by *Paenibacillus* sp. strain UY79 exerts antagonistic activity against different fungi and oomycetes. Antagonism was analyzed by the two base plate method. Production of volatile compounds with antimicrobial activity was evidenced as a growth inhibition of the fungi/oomycetes faced to strain UY79 (bottom plates) in comparison with the fungal growth not faced to the bacterium (upper plates). Fungi/oomycetes analyzed were: *Botrytis cinerea* A1 (**a**), *Fusarium graminearum* S127 (**b**), *Fusarium oxysporum* J38 (**c**), *Fusarium semitectum* J41 (**d**), Fusarium *verticillioides* A71 (**e**), *Macrophomina phaseolina* J431 (**f**), *Phomopsis longicolla* J429 (**g**), *Phytophthora sojae* Ps25 (**h**), *Pythium ultimum* Py03 (**i**), *Rhizoctonia solani* Rz01 (**j**), *Sclerotium rolfsii* 1948 (**k**) and *Trichoderma atroviride* 1607 (**l**). Assays were performed on PDA medium with the exception of *P. sojae* Ps25 that was grown on V8 agar medium. Experiment was done in triplicate.**A**) Photographs of one representative plate per treatment.**B**) Percentage of mycelial growth inhibition (mean values ± standard deviation).

As hydrogen cyanide (HCN) is a VC with well-known antimicrobial activity (49) and its production has been reported in some species of *Paenibacillus* (50), we assessed the ability of strain UY79 to produce this compound. According to the picrate-filter paper method, no production of HCN was observed in either PDA or TSA medium, neither with nor without the addition of glycine (data not shown). These results suggest that antimicrobial VCs other than HCN are being produced by strain UY79.

Other putative antagonistic traits, such as the production of cell wall degrading enzymes, siderophore production and proteolytic activity were assessed. As shown in Fig. 5, strain UY79 exhibited cellulase (Fig. 5A), β-glucosidase (Fig. 5B) xylanase (Fig. 5C), and protease (Fig. 5D) activity. Remarkably, strain UY79 displayed better cellulase and β-glucosidase activity than the positive controls, which are laboratory strains harboring a plasmid containing a constitutively expressed cellulase or β-glucosidase, respectively. Moreover, xylanase activity was evidenced even without the need of congo red staining, suggesting high enzymatic activity. These results show that strain UY79 harbors a suit of cell wall degrading enzymes which might be responsible for the antagonism mediated by diffusible compounds. Moreover, as depicted in Fig. 5D, strain UY79 was able to produce siderophores as observed in CAS medium.

**Fig 5.**
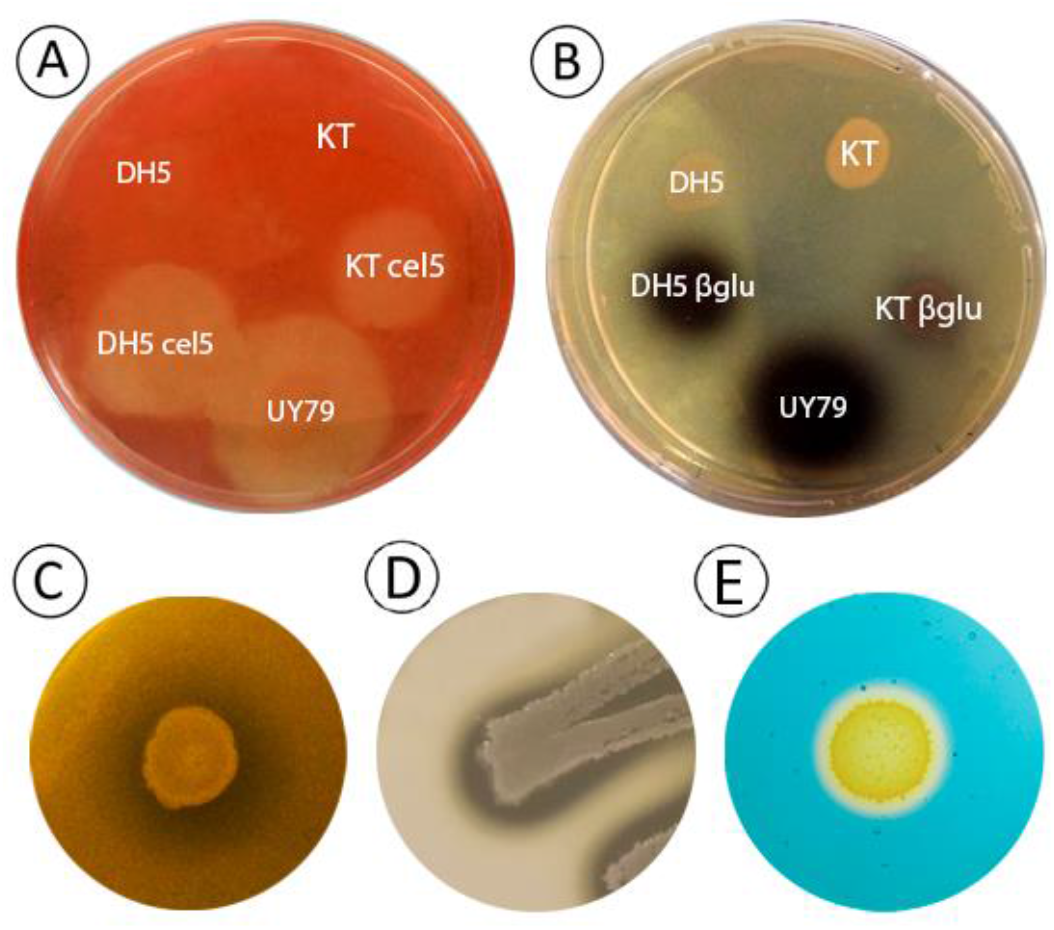
*Paenibacillus* sp. strain UY79 exhibits cellulase, β-glucosidase, xylanase and protease activity, as well as siderophore production.**A)** Cellulase activity was detected as a clear halo around colonies grown on carboxymethyl cellulose (CMC) containing medium after staining with congo red. *E. coli* DH5α cel5 (DH5 cel5) and *P. putida* KT2440 cel5 (KT cel5) were used as positive controls while *E. coli* DH5α (DH5) and *P. putida* KT2440 (KT) as negative controls.**B)** Beta-glucosidase activity was detected as a dark halo around colonies grown on esculin containing medium. *E. coli* DH5α β glu (DH5 βglu) and *P. putida* KT2440 β glu (KT βglu) were used as positive controls while *E. coli* DH5α (DH5) and *P. putida* KT2440 (KT) as negative controls.**C)** Xylanase activity was detected as a clear halo around *Paenibacillus* sp. UY79 colonies on xylan containing medium.**D)** Proteolytic activity was detected as a clear halo around *Paenibacillus* sp. UY79 colonies on skim milk containing medium.**E)** Siderophore production was detected in CAS medium as an orange halo around bacterial colony.

### Putative plant-growth promotion traits were not detected

With the aim to evaluate if strain UY79 displayed direct plant-growth promotion (PGP), we assessed the presence of *nifH* gene, production of IAA-like compounds and solubilization of phosphate compounds. No *nifH* gene was detected by PCR (data not shown) and neither IAA-like compounds nor phosphate solubilization compounds could be detected by *in vivo* analysis (Fig. S2). These results suggest that main PGP mechanisms are not present in strain UY79, and therefore it is probably not capable of exerting PGP.

### *Paenibacillus* sp. UY79 did not affect alfalfa (*Medicago sativa* L.) plant growth promotion by rhizobia

Keeping in mind the potential use of strain UY79 as a biopesticide for diverse crops, and taking into account that alfalfa is the most important legume crop in cultivated area after soybean (51), we assessed the effect of UY79 on alfalfa plant growth promotion exerted by rhizobia. As shown in Fig. 6A and 6B, inoculation of alfalfa plants with strain UY79 did not enhance plant dry weight, suggesting this strain has no PGP effect, which is in agreement with the aforementioned results, where no common PGP traits (P-solubilization, IAA production, *nifH* gene) were detected (Fig. S3). More importantly, plant growth promotion exerted by rhizobia (evaluated as shoot dry weight) (Fig. 6A and 6B) and number of nodules per plant (Fig. 6C and 6D), were not significantly affected by UY79 strain.

**Fig 6.**
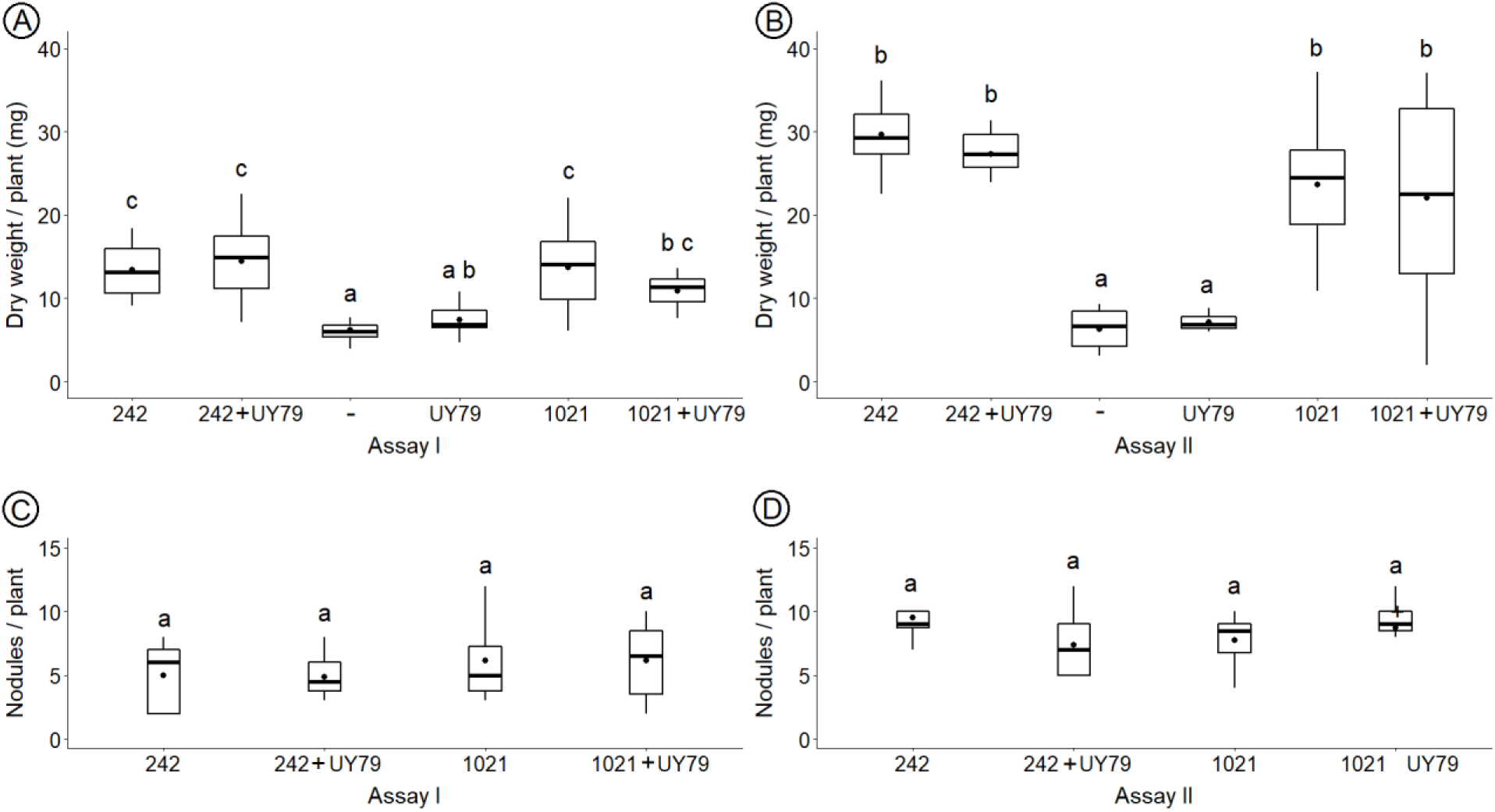
*Paenibacillus* sp. UY79 does not affect plant growth promotion exerted by rhizobia. Evaluation of plant-growth (**A** and **B**) and nodule number per plant (**C** and **D**) in response to inoculation is shown in the figure. Results of assay I are depicted in **A** and **C** and of assay II in **B** and **D**. *Medicago sativa* cv crioula plants were grown in N-free medium. *E. meliloti* 242, *E. meliloti* 1021 and *Paenibacillus* sp. UY79 strains were used to inoculate alfalfa seedlings separately or as a mixture (1:1) of the rhizobium strain with *Paenibacillus* sp. UY79 (242+UY79 and 1021+UY79, respectively). A negative control (-) without bacteria was also included. Data from each graph were independently analyzed as described in materials and methods section, and different letters in the same graph indicate significant differences.

### Assessment of antibacterial activity

Considering the potential use of the strain UY79 as a biological control agent, and therefore its possible release into the environment, we investigated its capability to coexist with or to inhibit diverse soil and plant associated bacteria. As shown in Fig. S3, antibiosis phenotype was diverse and depended on the target strain, as well as on the medium where the assay was performed. Strain UY79 exerted no growth inhibition and co-existed with rhizobia strains *E. meliloti* 1021, *R. tropici* CIAT 899 and *Paraburkholderia* sp. UYPR4.13, while for *B. elkanii* U-1301 and U-1302, type I inhibition was observed. Interestingly, in the case of *C. necator* UYPR2.512 a medium-dependent phenotype was observed, as UY79 exerted no inhibition in TY and type II inhibition on TSA. This medium-dependent phenotype was also observed for gram-positive *B. subtilis* ATCC6633, which was highly inhibited in TY (type IV inhibition) but was not affected in TSA, where both strains co-existed. When other PGP-bacteria were assessed, strain UY79 exerted type III inhibition in *A. brasilense* Sp7 and *Streptomyces* sp. UYFA156, while in the case of *Pseudomonas*, strain UY79 was not capable of growing. Lastly, in the case of the plant pathogen *E. caratovora* SCC3193 no inhibition by strain UY79 was observed in the condition assayed. In order to investigate if antibacterial compounds were present in the supernatant of a UY79 culture, we performed a similar assay but with a drop of a 45µm-filtered supernatant of UY79 over the confluent lawn of the target strain. No inhibition was detected with this approach (data not shown). This result suggests that other factors are required to generate the inhibitory phenotype such as a physical contact between strain UY79 and target strains, a bacterial biofilm formation. Another plausible explanation is that antibacterial compounds present in the supernatant did not reach a concentration sufficient to exert inhibition.

### Genome mining of genes putatively involved in antimicrobial response or in PGP activity

In agreement with results obtained by *in vivo* analysis, we did not find a *nifH* homologous in the UY79 genome reinforcing the fact that strain UY79 would not able to fix nitrogen (Table S3). Moreover, auxin production genes were not detected in the genome and the only function putatively related to phosphate solubilization was alkaline phosphatase (ID= UY79_2289) (Table S3). These results, together with data obtained in the alfalfa plant assay, where no plant promotion was observed for plants inoculated with UY79 (Fig. 6), clearly indicate that strain UY79 is not a PGP-bacterium.

Further, we manually scored the RAST 2.0 annotation of the UY79 draft genome searching for functions involved in the biological control of phytopathogenic fungi and oomycetes. Several genes involved in the synthesis of hydrolytic enzymes were identified (Table S3), among them seven cellulases, three β-glucosidases, sixteen xylanases, two chitinases and one protease. We also found genes involved in the biosynthesis of the achromobactin-like siderophore class and also genes putatively involved in the internalization of bacillibactin, ferrichrome and other siderophores. Also, several genes of polyketide synthase modules and related proteins presumptively involved in biological control were identified. No genes for HCN synthesis were identified in agreement with results obtained by *in vitro* assay.

In order to get closer to the possible compounds with antimicrobial activity produced by *Paenibacillus* sp. strain UY79, we used the antiSMASH 5.2.0 web server. Six non-ribosomal peptide synthetase or polyketide clusters (NRPS/PKS) were identified. NRPS are multidomain mega-enzymes which synthesize peptides with a vast range of bioactivities. Among the six detected clusters identified by antiSMASH three showed more than 40 % similarity with known NRPS reported to be involved in the synthesis of: Tridecaptin (100 % cluster similarity), Fusarisadin B (100% cluster similarity) and Tridecaptin (40 % cluster similarity). The remaining three NRPS did not show similarity with known clusters. We also found one ribosomally synthesized and post-translationally modified peptide (RiPP) cluster putatively involved in the synthesis of the lasso peptide type identified as a Paeninodin, with a 40 % cluster similarity.

By using PROPHAGE HUNTER bioinformatic tools, we identified eight prophage regions in the genome of strain UY79, five of them predicted as active and the remaining three as ambiguous. Meanwhile, PHASTER detected only five regions, two predicted as active and three as inactive (Table S3).

Interestingly we detected several CRISPR related regions: eighty-four CRISPR repeats and seventy-six CRISPR spacers, distributed among eight CRISPR arrays, and two CRISPR/CAS systems with the following protein arrangement: Cas3-Cas5-Csd1(Cas8c)-Csd2/Csh2(Cas7)-Cas4-Cas1-Cas2, corresponding for putative Type I-C system; and Cmr6-Cmr1-Cmr2-Cmr3-Cmr4-Cmr5-Cmr6, corresponding for putative Type III-B system (Table S3). Similar systems have also been described in *P. polymyxa* A18 (Type I-C and III-B), *Bacillus halodurans* (Type I-C) and *Pyrococcus furiosus* (Type III-B) (52, 53).

## DISCUSSION

In this study we report the isolation, identification, functional characterization and genome analysis of a *Paenibacillus* sp. strain UY79, recovered from an *A. villosa* nodule. It has been previously reported that bacteria belonging to the genus *Paenibacillus* are members of the root-nodule microbiome (13, 14, 54). The function that these bacteria exert in the nodule is not completely understood but, of particular relevance to the present study, is the fact that many *Paenibacillus* species have been reported to have antagonistic activities against different phytopathogenic fungi/oomycetes (55, 56). Moreover, diverse commercial biocontrol agents are based on *Paenibacillus* species (55). These observations prompted us to investigate *Paenibacillus* sp. strain UY79 potential as a biocontrol agent.

According to ANI analysis of twenty-two *Paenibacillus* genomes publicly available (Fig. 1), 16S *rRNA* phylogeny and MLSA, we conclude that strain UY79 is a new *Paenibacillus* species that belongs to the *P. polymyxa* complex (Fig. 1 and Fig. S1). This complex harbor six species of *Paenibacillus* (*P. kribbensis, P. peoriae, P. brasilensis, P. polymyxa, P. ottowii* and *P. terrae*) and until very recently no representatives were isolated from root nodules. In 2021, Ali *et al*. (45) reported the isolation of *P. peoriae* from nodules of *Robinia pseudoacacia* and *Dendrolobium triangulare*, and of *P. kribbensis* from *Ormosia semicastrata* nodules; therefore, so far as we know, *Paenibacillus* sp. UY79 is the third representative of the *P. polymyxa* complex to be isolated from root nodules. Further investigations should be undertaken in order to establish which traits contribute for *Paenibacillus* species to be part of the *P. polymyxa* complex, and why *Paenibacillus* species are frequent members of the legume nodule microbiota. Comparative genomic studies between nodule-inhabiting and other *Paenibacillus* strains could help in the near future for the comprehension of this behavior.

Regarding functional characterization, we can conclude that strain UY79 has a striking antagonism ability. By production of diffusible compounds, strain UY79 was capable to inhibit the growth of all the fungi (including relevant phytopathogenic fungi) and the two phytopathogenic oomycetes analyzed (Fig. 2). Interestingly, antagonistic activity was present in the cell-free supernatant of a UY79 culture grown for up to 96 h (Fig. 3). This is a promising trait in view of a possible application of these inhibitory compounds in the field as pathogen suppressors.

According to genome mining, a cluster presumptively involved in the synthesis of a fusaricidin B-like compound was found (Table S3). Fusaricidins are NRPs, consisting of a cyclic lipopeptide with six amino acids and an unusual fatty acid chain of 15-guanidino-3-hydroxypentadecanoic acid (57). There are four types of known fusaricidin (fusaricidin A-D), all of which display a general peptide sequence of L-Thr–X1–X2–D-allo-Thr–X3–D-Ala. Fusaricidin exhibit antagonistic activity against diverse fungi as well as against gram-negative bacteria (58). Although a NRPS presumptively involved in the synthesis of fusaricidin B was detected by using the antiSMASH bioinformatic tool, the predicted peptide sequence was Ser-D-Val-Ser-Ser-D-Asn-Ala, which is different to that reported for known fusaricidins. Therefore, with the data available to date, we cannot establish if strain UY79 encodes a new fusaricidin or a different compound with similar molecular structure to that of fusaricidins.

Genome mining with antiSMASH bioinformatic tool also predicted other five NRPS as well as one ribosomally-synthesized and post-translationally modified peptides (RiPPs). Two NRPS clusters are probably involved in the synthesis of tridecaptin-like compounds (Table S3). Tridecaptins consist of linear acylated tridecapeptides that exhibit a strong selective antibacterial activity against gram-negative bacteria and moderate activity against gram-positive bacteria (59, 60). They are produced by *Bacillus* and *Paenibacillus* species, including by strains of the *P. polymyxa* complex (61, 62).

Putative protein encoded by UY79_3797 presented homology with the lasso peptide paeninodin. Lasso peptides are RiPPs with a unique structure comprising 16-21 residues, produced by clusters of unique and well conserved organization (63). Interestingly, it has been found that some lasso peptides possess antimicrobial and antiviral activities.

Results obtained by *in vitro* antibiosis assays against various soil bacteria, showed that strain UY79 was indeed capable of inhibiting the growth of some strains of gram-positive and gram-negative bacteria (Fig. S3), which is in agreement with the predictions made by genome mining.

Other mechanisms involved in the protection of plant diseases caused by phytopathogens are based on the production of hydrolytic enzymes. Enzymes able to degrade the cell wall of fungi or oomycetes (chitinases, cellulases, β-1,3-glucanases) antagonize the mycelial growth, while xylanases may act as elicitors of plant systemic resistance, enhancing plant resistance to pathogens (64, 65). Moreover, cellulases and β-glycosidases may be implicated in facilitating plant tissue colonization enabling bacterial endophytic lifestyle (55). By functional analysis we found that strain UY79 displayed β-glucosidase, cellulase and xylanase activities (Fig. 5), and by genome mining we identified several genes putatively encoding these enzymes (Table S3), which supports the activities observed *in vitro*. However, it should be noted that the presence of these hydrolytic activities or encoding genes does not indicate that they are in fact involved in biological control. For instance, Ali *et al*. (45), recently found that β-1,3-glucanase and chitinase activity exhibited by some *Paenibacillus* sp. strains were not involved in their antifungal activity.

Different roles have been assigned to siderophores produced by plant-associated bacteria. They may be responsible for pathogenicity, such as the siderophore produced by *Erwinia chrysanthemi*, or may promote plant growth through iron solubilization or by controlling phytopathogens (66-68). We currently do not have enough information to ascertain the putative role of siderophore(s) produced by strain UY79 in biocontrol; nonetheless, considering that the antagonistic assays performed in this work were done in iron sufficient PDA medium, we can speculate that siderophore(s) were not being produced and therefore did not contribute to the observed antifungal activity.

Strain UY79 was also capable of inhibiting the growth of most of the fungi/oomycetes assessed by the production of volatile compounds (VCs) (Fig. 4), being the only exception the plant growth promoter *T. atroviride* 1607. This effect was not mediated by HCN as no production of this VC could be detected, and no genes for HCN production could be found (Table S3). Investigations of the volatilome (including organic as well as inorganic VCs) produced by bacteria is a topic that has aroused great interest. Using VCs for biocontrol of plant pathogens present some advantages, as they may induce systemic resistance in plants, can be used in conditions where physical contact between the pathogen and the BCA is not possible, and leaves less residuals in the environment once applied (56). Moreover, volatile compounds may participate in a plethora of interaction intra and interkingdom (69) which makes these compounds a fascinating subject of research.

It is interesting to highlight that by genome mining, we found the presence of different systems putatively involved in antibiosis and bacteriophage production, as well as several regions involved in the CRISPR/Cas adaptative antiviral immune response. These results together with the ability to produce diverse diffusible and volatile compounds against fungi/oomycetes suggest that strain UY79 could either coexist or interfere with various soil and plant-associated microorganisms, perhaps modulating the microbiota associated with plants, as has been described for other nodule-inhabiting *Paenibacillus* strains (14, 70).

Many *Paenibacillus* species have been reported as plant growth promoters, not only indirectly by their antagonism effect, but also by direct mechanisms (45). Results obtained in this work indicate that strain UY79 might not have the ability to improve plant growth directly, as it lacks many of the main mechanisms previously reported as important for this effect (nitrogen fixation, auxin production, and phosphate solubilization) (Fig. S3, Table S3). Moreover, no plant growth promotion was evidenced in alfalfa plants when inoculated with strain UY79 (Fig. 6). In that sense, Agaras *et al*. (71), after screening multiple traits putatively involved in biocontrol and direct plant growth promotion in *Pseudomonas*, found that “top fungal antagonistic potential together with the highest direct plant growth promotion potential rarely converge in a single *Pseudomonas* isolate”. A similar conclusion was achieved by Ali *et al*. (45) after performing a genome mining analysis in strains within the *P. polymyxa* complex.

Despite the fact that strain UY79 exert no promotion of alfalfa growth in gnotobiotic conditions, it is important to highlight that inoculation with strain UY79 did not interfere with plant growth promotion exerted by rhizobia (Fig. 6). Some chemical pesticides have probed to affect the rhizobia-legume symbiosis (72, 73), therefore, the fact that strain UY79 did not interfere with symbiosis or biological nitrogen fixation exerted by symbionts, are additional advantages when thinking of its use as a biological control agent. Nonetheless, the effect should be analyzed for each particular legume-rhizobia association.

In conclusion, we identify a new nodule-inhabiting *Paenibacillus* species that belongs to the *P. polymyxa* group. The wide spectrum of its antagonistic effect, together with the diversity of mechanisms involved (diffusible and volatile compounds, hydrolytic enzymes, iron scavenging), make *Paenibacillus* sp. UY79 a promising biocontrol agent.

